# ATP Mediates Phase Separation of Disordered Basic Proteins by Bridging Intermolecular Interaction Networks

**DOI:** 10.1101/2023.08.20.554035

**Authors:** Divya Kota, Ramesh Prasad, Huan-Xiang Zhou

## Abstract

ATP is an abundant molecule with crucial cellular roles as the energy currency and a building block of nucleic acids and for protein phosphorylation. Here we show that ATP mediates the phase separation of basic intrinsically disordered proteins (bIDPs). In the resulting condensates, ATP is highly concentrated (apparent partition coefficients at 200-5000) and serves as bridges between bIDP chains. These liquid-like droplets have some of the lowest interfacial tension (∼25 pN/μm) but high zero-shear viscosities (1-15 Pa s) due to the bridged protein networks, and yet their fusion has some of the highest speeds (∼1 μm/ms). The rapid fusion manifests extreme shear thinning, where the apparent viscosity is lower than zero-shear viscosity by over 100-fold, made possible by fast reformation of the ATP bridges. At still higher concentrations, ATP does not dissolve bIDP droplets but results in aggregates and fibrils.

## Introduction

ATP is present at high levels (2 -12 mM) in all cells and plays several crucial roles. It is best known as the energy currency, backed by an out-of-equilibrium state with its hydrolysis product ADP at levels far below equilibrium. It is a building block of DNA and RNA, nucleic acids that are the basis for the storage of genetic information and its translation to proteins. ATP is also the substrate molecule for an important class of protein posttranslational modification, i.e., phosphorylation. Recently a new role of ATP was identified, namely as a hydrotrope ^1^, which denotes small molecules that do not aggregate on their own but at high concentrations solubilize hydrophobic molecules. At physiological concentrations, ATP inhibited the phase separation of intrinsically disordered proteins (IDPs) including FUS and dissolved preformed FUS aggregates. The present study targets the effects of ATP on a different type of IDPs, namely those that are enriched in basic residues (arginine and lysine) and referred to as bIDPs.

Phase separation of bIDP-nucleic acid mixtures has been reported, many times for RNA ^2-20^ but also for DNA ^21^. Basic residues on the IDPs are important drivers of phase separation, forming electrostatic interactions with phosphate groups of the nucleotides and cation-π interactions with nucleobases. Between arginine and lysine, the former forms stronger interactions with RNA ^15^, in particular resulting in much higher viscosity ^16^ or reduced fluidity ^10, 11, 15^. At high concentrations, RNA suppresses phase separation ^6, 8, 9, 13^, as RNA molecules present in the droplet phase start to experience electrostatic repulsion among themselves. Similar results are obtained when RNA is replaced by acidic polymers, including heparin ^22^ and hyaluronic acid ^23^.

As a building block of nucleic acids, ATP should form similar interactions as RNA and DNA with basic residues of IDPs and therefore can be expected to mediate phase separation of bIDPs. Fisher et al. ^16^ demonstrated that poly-arginine (pR) and poly-lysine (pK) forms droplets when mixed with the nucleotide UTP as well as the polymerized form poly-uridine. Droplet formation of pK mixed ATP was shown at 30 mM ATP ^24^ and at physiological concentrations ^25^. Nobeyama et al. ^26^ reported the phase diagram of mixtures of pK with ATP at very high concentrations (50 to 500 mM). Dec et al. ^27^ found that pK with an N-terminal tag when mixed with ATP initially formed droplets, which then converted into fibrils. Whereas Patel et al. ^1^ reported inhibition of FUS phase separation by 8 mM ATP, Kang et al. ^28^ found that ATP at concentrations up to 1 mM actually promoted the FUS phase separation. Likewise, Dang et al. ^29^ showed that ATP up to 1.5 mM promoted the phase separation of the TDP-43 prion-like domain but the promotional effect diminishes with further increase in ATP concentration, reminiscent of the biphasic regulatory effects of RNA. Biphasic effects of ATP on the phase separation of the disordered C-terminal region of CAPRIN1 and the condensation of the folded basic protein lysozyme were also reported ^30, 31^. This effect was recapitulated by coarse-grained molecular dynamics (MD) simulations ^32^.

The interactions of ATP with proteins have been probed by various techniques. ^1^H-^15^N NMR HSQC spectra of FUS at increasing ATP concentrations showed that ATP, similar to single-stranded DNA, binds to arginine and lysine residues, but not to its aromatic-rich prion-like domain ^13, 28^. Similarly, in the TDP-43 prion-like domain, the residues with significant shifts in HSQC peaks upon titrating ATP centered around three arginines ^29^. After mutating the arginines to lysines, the peak shifts saturated at higher ATP concentrations, indicating lower affinities of ATP for lysine than for arginine. ATP also appears to bind to arginines in the cold inducible RNA binding protein ^20^. In another study based on ^1^H-^15^N HSQC, ATP binding was detected for basic and hydrophobic residues on two folded proteins but predominantly for an acidic region in the IDP α-synuclein. For lysozyme, ^13^C HSQC identified six ATP binding sites, all of which contain arginine but only one contains lysine, suggesting a strong preference of ATP for arginine over lysine ^31^. Three of these sites were captured by a lysozyme-ATP crystal structure. Interestingly, in the crystal lattice, the ATP molecules bridge between lysozyme molecules. Recently, NMR-based measurement of the surface electrostatic potential of CAPRIN1 was done at increasing concentrations of ATP. ATP gradually decreases the positive electrostatic potential of this bIDP. Upon phase separation, ATP neutralizes the electrostatic potential of the protein in the dense phase ^30^. In all-atom MD simulations of folded proteins including lysozyme, ATP molecules assemble into clusters mediated by cations; Na^+^ is as effective as Mg^2+^ in playing this role ^33^. ATP can in fact self-assemble into clusters, likely mediated by cations ^1, 34, 35^.

Here we used brightfield, transmission electron microscopy (TEM), and confocal imaging, optical tweezers (OT), and all-atom MD simulations to characterize the thermodynamic and material properties of bIDP-ATP condensates. Two bIDPs were studied (Fig. 1a): pK and the arginine-rich protein protamine (PM). Both PM-ATP and pK-ATP mixtures form droplets at physiologically relevant concentrations. The droplets have very unusual material properties, including fast fusion speed, low interfacial tension but high viscosity, leading to extreme shear thinning during fusion. Underlying these material properties, our MD simulations revealed that, in the dense phase, ATP molecules bridge between PM chains, with their phosphate groups forming salt bridges with multiple arginines and adenines forming multi-layered π - π and cation-π stacks.

**Fig. 1.**
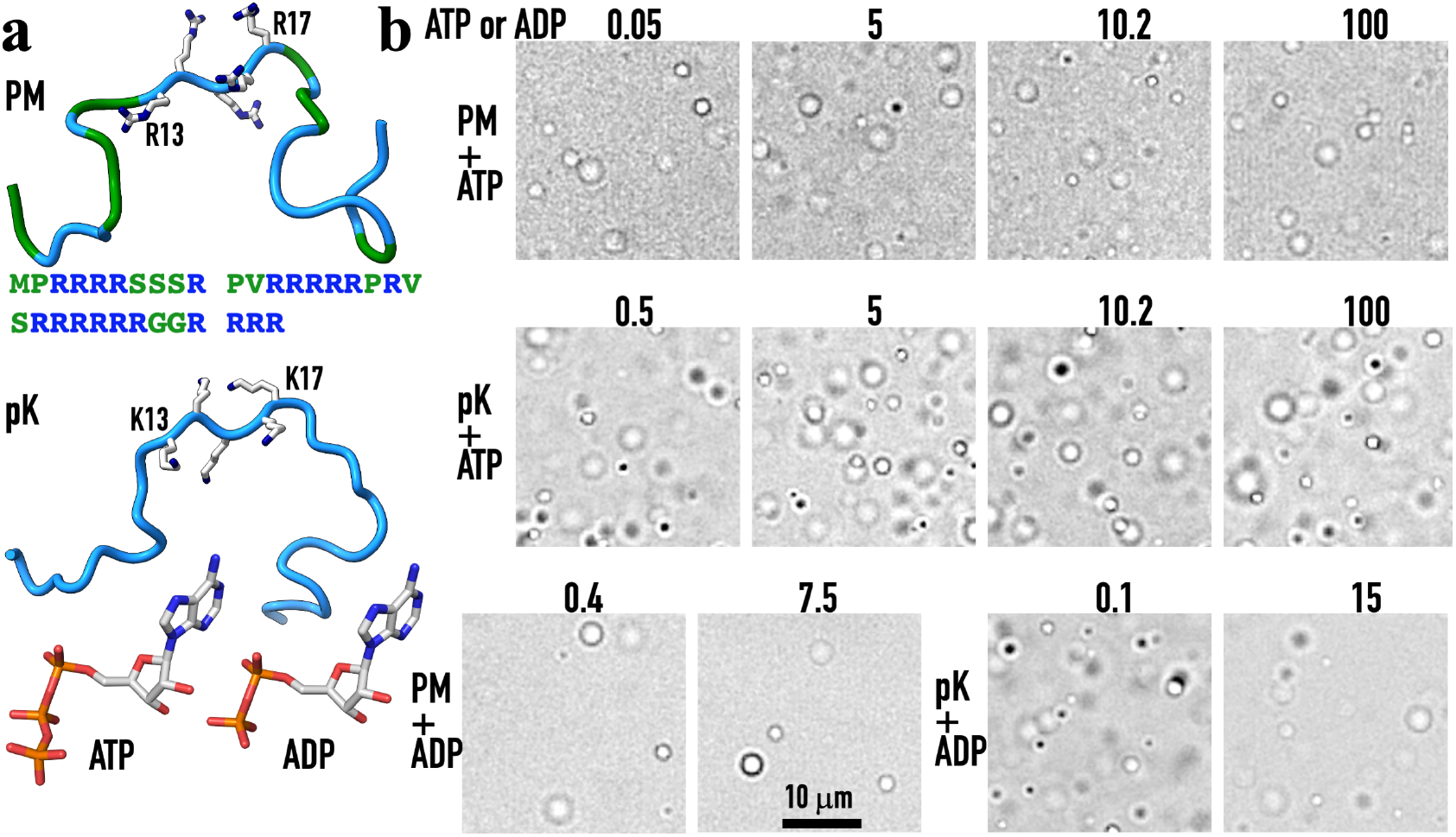
Droplet formation by mixing bIDPs with either ATP or ADP. (a) The bIDPs studied and ATP and ADP molecules. The sequence of PM is shown with 21 arginines in blue and other 12 residues in green; the PM chain is shown with backbone in colors matching those of the sequence and sidechains of R13 to R17 in stick representation. The pK chain is also shown with the sidechains of K13-K17 in stick representation. (b) Brightfield images of droplets formed by mixing 10 μM PM or pK with ATP or ADP at the indicated concentration (in mM).

## Results

To highlight the effects of ATP exerted via intermolecular interactions on the phase separation of bIDPs, we excluded from our samples Mg^2+^, which is typically required for enzymatic conversion of ATP. The buffer was 10 mM imidazole pH 7. Experiments were done both without added salt and with 150 mM KCl. Even without the added salt, the samples still contained Na^+^ from NaOH used for adjusting pH of the ATP stock solution and Cl^−^ from HCl used for adjusting the pH of the buffer.

### bIDPs form condensates over a wide range of ATP concentrations

At 10 μM PM, droplets are observed over a very wide range of ATP concentrations, from 0.05 to 100 mM (Fig. 1b, first row). As the ATP concentration is further increased, PM forms first amorphous aggregates (200 mM ATP), then fibril-like aggregates (500 mM ATP), and finally fibrils (750 mM ATP) (Fig. S1a). At pH 7, PM carries a net charge of +21*e* (Fig. 1a) whereas ATP carries a net charge of –4*e*. So charge neutralization would require a 21/4 = 5.25-fold excess of ATP over PM, which for 10 μM PM corresponds to 0.0525 mM ATP. The latter value corresponds to the start of the ATP concentration range for droplet formation. So apparently an unneutralized positive charge from the bIDP chains can lead to dissolution of droplets. On the other hand, droplets can withstand an enormous level of unneutralized negative charge from the ATP molecules. Indeed, in droplets formed at 10 μM PM and 100 mM ATP, the ATP concentration is approximately 2,000-fold higher than required for charge neutralization. In PM fibrils, the excess ATP level even reaches 15,000-fold of the neutralization concentration.

The second bIDP, pK, behaves in a similar way as PM in ATP-mediated phase separation. The range of ATP concentrations over which 10 μM pK forms droplets goes from 0.5 to 100 mM (Fig. 1b, second row). Again, at higher ATP concentrations, aggregates and fibrils are formed. Our pK has a molecular weight range of 15-30 kDa, corresponding to an average chain length of 200 lysine residues and thus a charge of +200*e*. The ATP concentration required for neutralizing 10 μM pK is 0.5 mM, which also coincides with the start of the ATP concentration range for droplet formation. The high end of the ATP concentration range for droplet formation is approximately 200-fold higher than required for charge neutralization.

ADP, the hydrolysis product of ATP, can also mediate the phase separation of bIDPs, but over a much narrower concentration range. At 10 μM PM, droplets are formed over 0.4 to 7.5 mM of ADP (Fig. 1b, third row left), compared with a concentration range of 0.05 to 100 for ATP. For 10 μM pK, the minimum ADP concentration for droplet formation is 0.1 mM, a little lower than the minimum of 0.5 mM ATP (Fig. 1b, third row right). However, at the opposite end, the maximum ADP concentration for pK droplet formation is 15 mM, of the same order of magnitude as in the case of PM droplets. Therefore, bIDP droplets can withstand an enormous level of unneutralized negative charge from the ATP molecules, but this ability is much curtailed for ADP.

We present the phase diagrams for PM and pK paired with ATP and ADP in Fig. 2, showing concentration regions where condensates are formed. The left boundary of the droplet-forming region approximately follows the line of charge neutralization, at which the nucleotide concentration is exactly what is required for neutralizing the charges on the bIDP chains at a given concentration. This line is followed most closely by pK-ATP droplets; PM-ADP and pK-ADP droplets have some incursions to the left of the line at PM above 50 μM and pK above 5 μM. For PM-ATP droplets, the incursions get deeper and deeper as the PM concentration is increased.

**Fig. 2.**
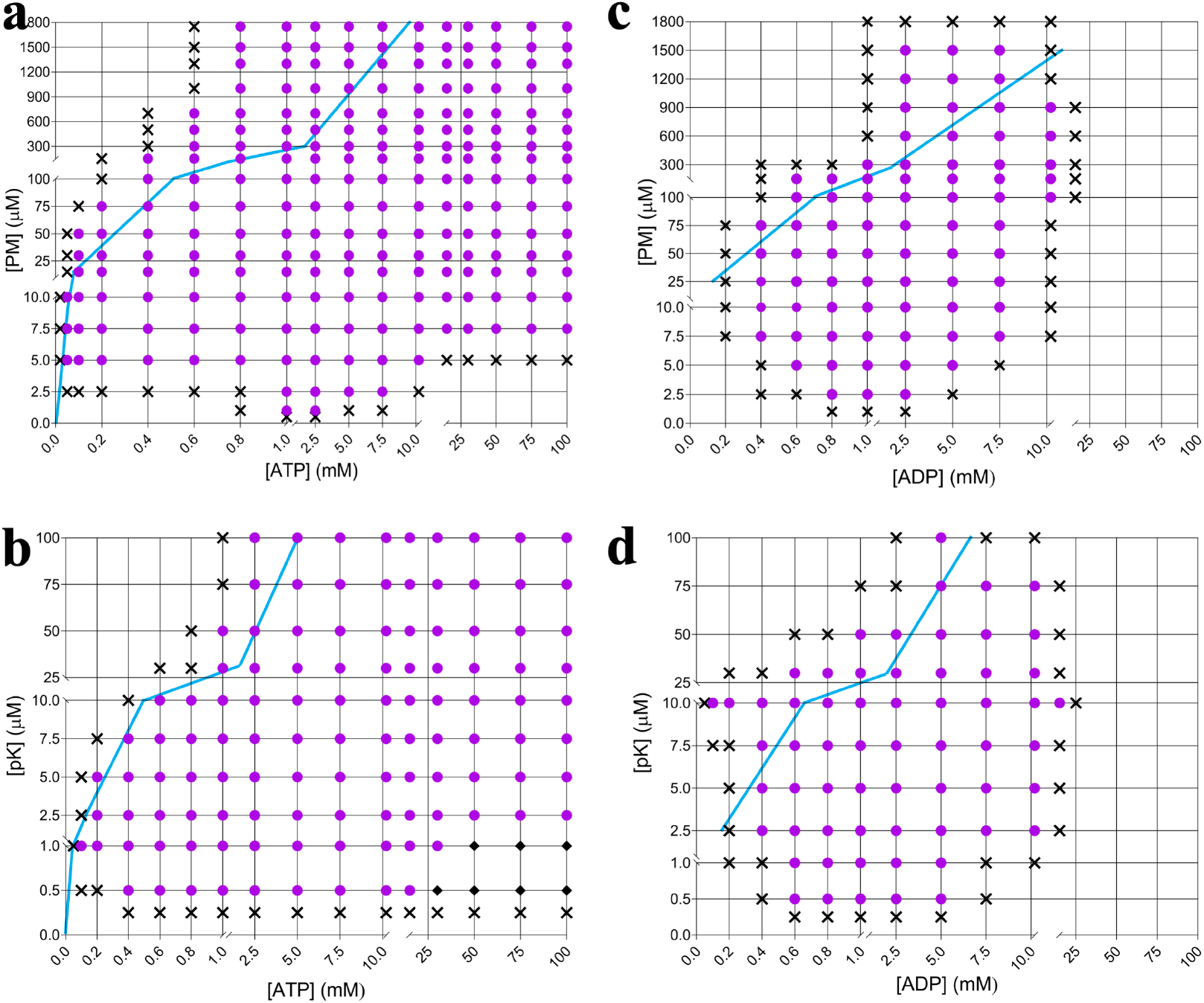
Phase diagrams of bIDP-nucleotide mixtures. (a) PM mixed with ATP. (b) pK mixed with ATP. (c) PM mixed with ADP. (d) pK mixed with ADP. Magenta circles indicate droplet formation; black diamonds indicate aggregate formation; crosses indicate homogeneous solution. Blue lines are drawn at concentration ratios producing charge neutralization. All phase diagrams are from observations on freshly mixed samples.

Next let us look at the bottom, or minimum bIDP concentration, of the droplet-forming region. For PM-ATP droplets, this minimum is 1.0 μM PM, at which droplets are observed over a narrow range of ATP concentrations, from 1.0 to 2.5 mM (Fig. 2a). As the PM concentration is increased from 1.0 μM, the range of ATP concentrations that result in droplet formation widens rapidly at both ends; at 7.5 μM PM, the range of ATP concentrations in the droplet-forming region is as wide as seen at 10 μM PM, i.e., from 0.05 to 100 mM ATP. As already noted, a further increase in ATP concentrations leads to aggregation formation (Fig. S1a).

For pK-ATP droplet formation, the minimum pK concentration is 0.5 μM (Fig. 2b). Whereas no aggregates are observed for PM-ATP mixtures at the lowest PM concentrations (Fig. 2a), we already see aggregates at 0.5 μM pK and ≥ 30 mM ATP. However, at 2.5 μM or higher pK, only droplets, not aggregates, are observed for up to 100 mM ATP. For mixtures of both bIDPs with ADP, no aggregates are ever observed, and the highest ADP concentration at which droplets are formed is 10.2 mM for PM and 15 mM for pK (Fig. 2c, d).

We also investigated the effects of salt on the phase diagrams. The phase diagram of pK-ATP mixtures at 150 mM KCl is presented in Fig. S2a. Salt affects the phase diagram in two locations. The first is at low ATP concentrations, where the minimum ATP concentration for droplet formation increases from 0.1 mM without KCl to 0.8 mM with 150 mM KCl. At these low ATP concentrations, Cl^−^ can compete out some of the ATP molecules and thereby weaken the intermolecular interaction networks in droplets. This effect is reminiscent of the situation with macromolecular regulators acting as weak-attraction suppressors of phase separation ^22, 36^. The second location where 150 mM KCl affects the phase diagram is at high ATP concentrations, where it also narrows the range of ATP concentrations over which droplets are observed. Here KCl exerts this effect by lowering the ATP concentrations at which aggregates are formed. We also examined by transmission electron microscopy (TEM) samples identified by brightfield images as aggregates, confirming their classification as either amorphous or fibril-like (Fig. S1b). In the presence of 150 mM KCl, we now finally see an area where aggregates no longer form upon a further increase in ATP concentration (lower right corner in Fig. S2a).

A location where KCl does not affect the phase diagram is when pK is at the highest concentrations studied (≥ 75 μM). Here pK forms droplets over the same range of ATP concentrations (2.5 to 100 mM), whether 150 mM KCl is present (Figs. 2b and S2a). Still, bright-field images suggest that KCl reduces the number of droplets formed at both the low and high end of this ATP concentration range (Fig. S2b).

In the experimental studies below, we use 100 μM bIDP because this concentration produces plentiful droplets, thereby facilitating their characterizations.

### ATP is highly concentrated in bIDP condensates

To get a sense of ATP concentrations inside bIDP droplets, we mixed Alexa Fluor 647 ATP (hereafter ATP*, at 1 μM) with unlabeled ATP in preparing droplets. The intensities of ATP* fluorescence show enormous differences inside and outside both PM-ATP and pK-ATP droplets (Fig. 3a, b). By determining a standard curve (Fig. S3a), we converted the fluorescence intensities to ATP concentrations (assuming the same ATP-to-ATP* ratio as in the initial concentrations). For droplets formed by mixing 100 μM PM with either 5 or 10.2 mM ATP, the partition coefficient of ATP* is 5000; the partition coefficients are 400 and 230, respectively, for droplets formed by mixing 100 μM pK with either 5 or 10.2 mM ATP (Fig. 3c). These partition coefficients are in line with the above observation that ATP can sustain bIDP droplet formation at concentrations that far exceed that for charge neutralization. 150 mM KCl has a modest effect on ATP* partition coefficients, reducing those in droplets formed by mixing 100 μM pK with either 5 or 10.2 mM ATP to 300 and 180, respectively (Fig. S3b).

**Fig. 3.**
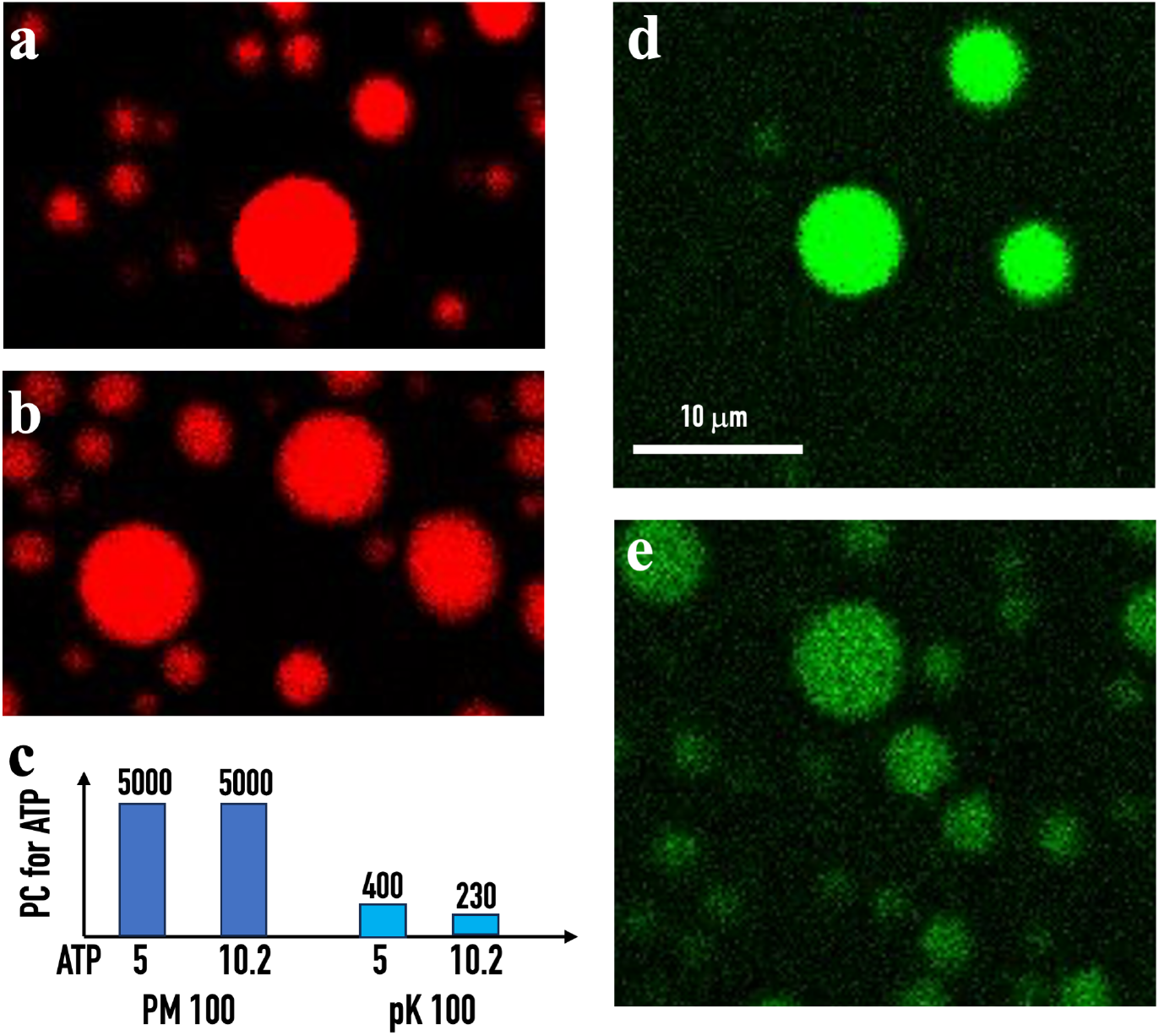
Partition of ATP and bIDP inside droplets by confocal microscopy. (a) Images of PM-ATP droplets, formed by mixing 100 μM PM with 5 mM ATP and labeled with 1 μM ATP*. (b) Corresponding images of pK-ATP droplets. (c) Partition coefficients (PCs) of ATP* in droplets formed by mixing 100 μM PM or pK and 5 or 10.2 mM ATP. (d) Images of PM-ATP droplets, formed by mixing 100 μM PM with 5 mM ATP and labeled with PM-FITC. (e) Corresponding images of pK-ATP droplets. The scale bar in (d) applies to all the confocal images.

We also labeled the bIDPs with FITC to get rough estimates for their concentration ratios inside and outside droplets. In droplets formed by mixing 100 μM PM with 5 mM ATP, PM-FITC fluorescence intensities are 20 times higher than in the dilute phase (Fig. 3d). In comparison, this ratio is 8 for the pK counterpart (Fig. 3e). As in many cases of fluorescently labeled proteins, we were not able to determine standard curves for the FITC-labeled bIDPs because we could not obtain them at sufficiently high concentrations. These intensity ratios likely underestimate the concentration ratios.

### bIDP-ATP droplets undergo rapid fusion

Fusion is a very important manifestation of the liquidity of biomolecular condensates. Fusion kinetics can be quantified by using dual-trap optical tweezers (OT) to trap two equal-sized droplets and direct them to fusion (Fig. 4a) ^37, 38^. We fit the fusion progress curve to a stretched exponential,

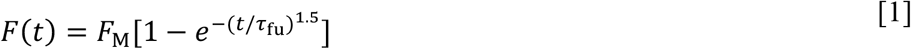

to obtain the fusion time τ_fu_ (Fig. 4b, c). As in previous studies ^6, 7, 37^, fusion time is proportional to the initial droplet radius (*R*) (Fig. 4d, e). The ratio τ_fu_*/R* is the inverse fusion speed. For droplets formed with 100 μM PM and 5 or 10.2 mM ATP, the inverse fusion speed is 11.9 ± 0.4 and 13.2 ± 0.8 ms/μm, respectively (Fig. 4f). These values are similar to those for droplets formed by oppositely charged IDPs ^37^. In comparison, the fusion speeds of pK-ATP droplets are increased by 5-10 fold, making them among the fastest observed for biomolecular condensates. Moreover, with increasing ATP concentration, τ_fu_*/R* increases slightly for PM-ATP droplets but exhibits a clear decreasing trend for pK-ATP droplets.

**Fig. 4.**
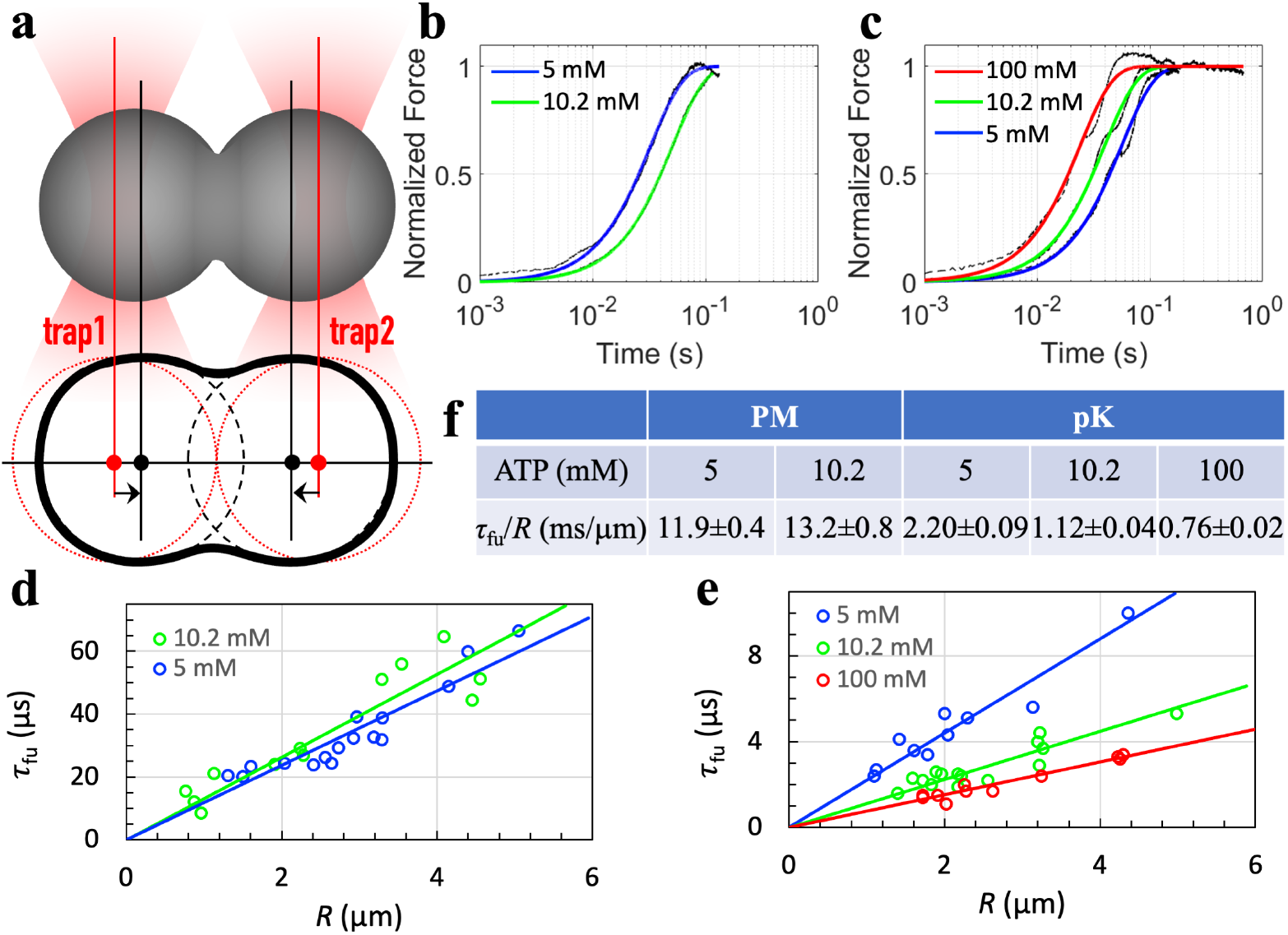
OT-directed droplet fusion. (a) Illustration of directed fusion. (b) Fusion progress curves of PM-ATP droplets. Black traces show raw data; colored curves are fits to Eq [1]. Droplets were formed by mixing 100 μM PM with ATP at the indicated concentration and grown to approximately 3 μm radius before fusion. (c) Corresponding results for pK-ATP droplets. (d) Fusion time of PM-ATP droplets as a function of initial droplet radius. Each circle represents a fusion event. Lines display the proportional relation between fusion time and droplet radius. (e) Corresponding results for pK-ATP droplets. (f) Inverse fusion speeds of bIDP-ATP droplets.

150 mM KCl increases the fusion speeds of both PM-ATP and pK-ATP droplets (Fig. S4). The fusion results at 150 mM KCl further confirm the opposite trends between PM-ATP and pK-ATP droplets in their changes in fusion speed with increasing ATP concentrations.

### bIDP-ATP droplets have low interfacial tension

Droplet fusion is driven by interfacial tension. Based on the fast fusion speeds of PM-ATP and pK-ATP droplets, we expected high interfacial tension. By using optically trapped beads to stretch droplets (Fig. 5a, b) ^39, 40^, we measured the interfacial tension of these droplets (Fig. 5c, d). To our surprise, the interfacial tension of PM-ATP droplets is only 24 pN/μm (Fig. 5e), among the lowest observed for biomolecular condensates. For pK-ATP droplets, the interfacial tension is 60 pN/μm at 5 mM ATP and progressively decreases to 27 pN/μm at 100 mM ATP. These low interfacial tension values are also obtained in the presence of 150 mM KCl (Fig. S5).

**Fig. 5.**
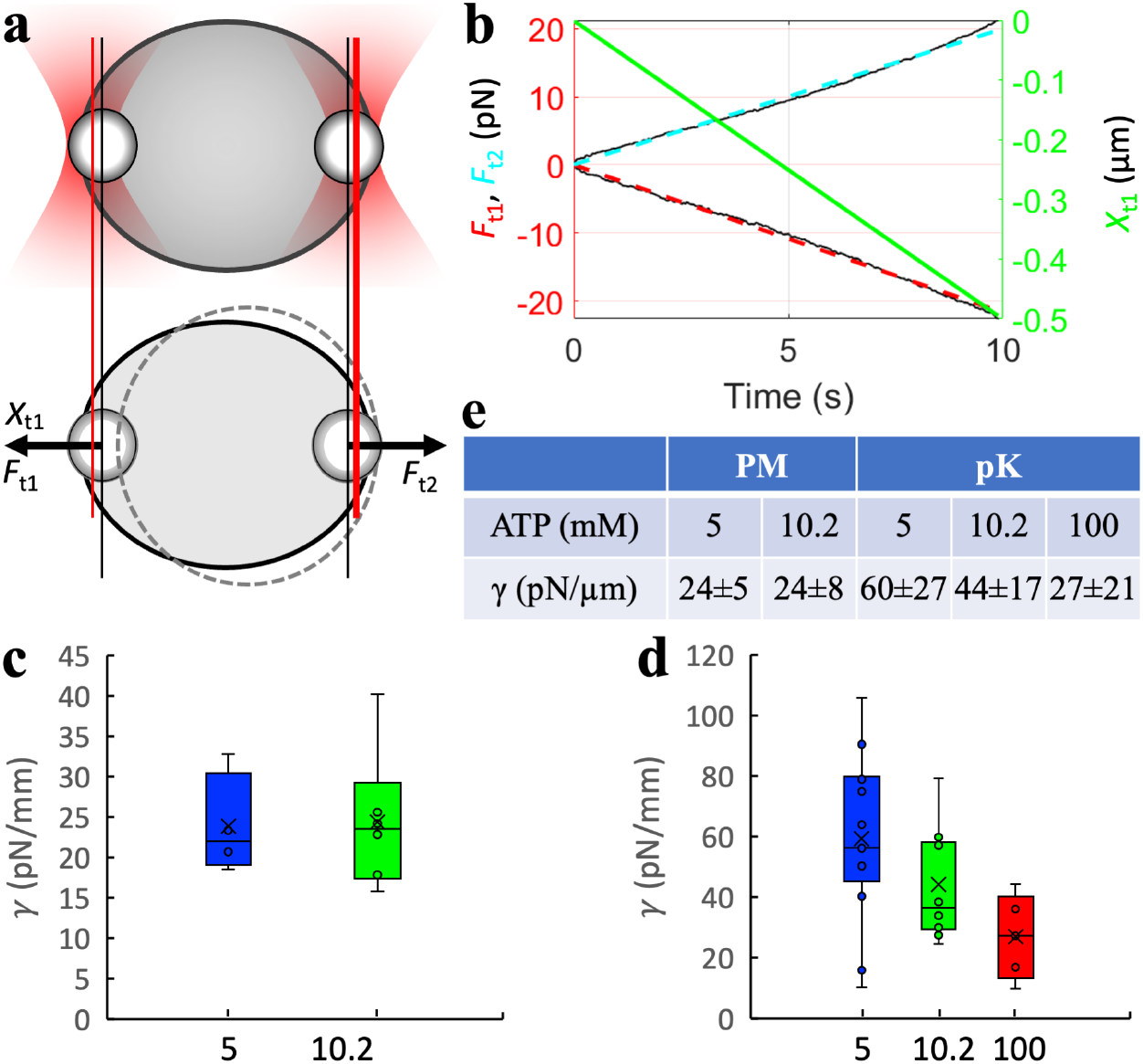
Interfacial tension of bIDP-ATP droplets. (a) Stretching of droplets by two optically trapped beads. (b) Trapping forces and extension measured on a droplet formed by mixing 100 μM pK with 5 mM ATP. (c) Interfacial tension of PM-ATP droplets (100 μM PM and ATP at indicated concentration in mM), displayed as a box plot. Circles represent values from individual droplets. The mean value is displayed as a cross. (d) Corresponding results for pK-ATP droplets. (e) Mean values of interfacial tension.

### bIDP-ATP condensates are highly viscous

For purely viscous droplets, which are formally known as Newtonian fluids, the inverse fusion speed is inversely proportional to interfacial tension (γ) and proportional to zero-shear viscosity (η) ^37^:

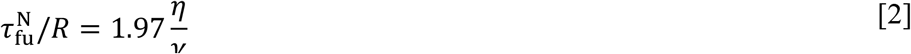

So conceivably the fast fusion speeds observed on bIDP-ATP droplets can still be explained by low viscosity. We have demonstrated that biomolecular condensates are viscoelastic rather than purely viscous, and the inverse fusion speed can deviate from the prediction for Newtonian fluids ^40^. By trapping a bead inside droplets and oscillating over a range of frequencies (*f* in Hertz or ω = 2π*f* in rad/s) (Fig. 6a), we determined the viscoelasticity of bIDP-ATP droplets. The amplitude of the corresponding trapping force and its phase shift from the oscillating bead position (Fig. 6b) allow for the determination of the elastic modulus *G*^*′*^(ω) and viscous modulus *G*″(ω). We fit the frequency dependences of *G*^*′*^(ω) and *G*″(ω) to the Jeffreys model (Fig. 6c):

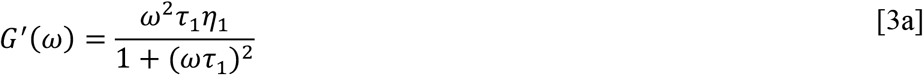

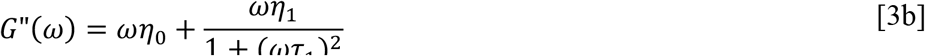

where η_0_ + η_1_ gives the zero-shear viscosity and τ_1_ is the shear relaxation time. The resulting values of η_1_, η_1_, and τ_1_ are collected in Fig. 6d.

**Fig. 6.**
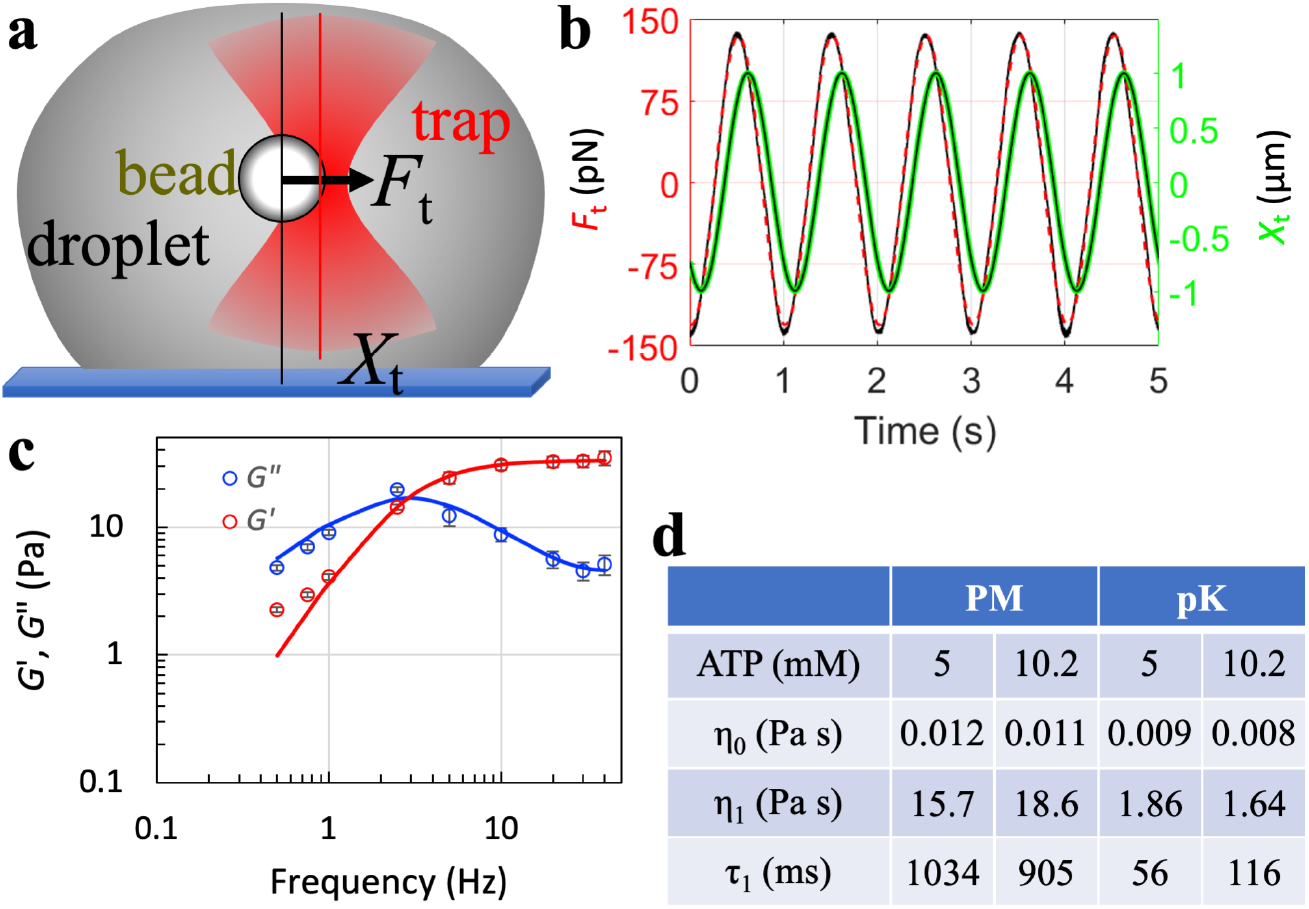
Viscoelasticity of bIDP-ATP droplets. (a) Oscillation of a trapped bead inside a droplet. (b) Time traces of bead position and trapping force, acquired in a droplet formed by mixing 100 pK with 5 mM ATP. (c) Elastic and viscous moduli of pK-ATP droplets (100 pK and 5 mM ATP). (d) Viscosities and shear relaxation times of bIDP-ATP droplets.

Both pK-ATP and PM-ATP droplets have high zero-shear viscosities. For pK-ATP droplets, η is 1.9 Pa s at 5 mM ATP and 1.6 Pa s at 10.2 mM ATP. These values are about 5-fold higher than those in droplets formed by oppositely charged IDPs ^40^. The viscosities of PM-ATP droplets are 10 times higher, pushing them beyond even that inside droplets formed by structured domains ^40^.

In short, the bIDP-ATP droplets have very unusual material properties, with some of the highest fusion speeds, some of the lowest interfacial tension, and some of the highest viscosities.

### ATP mediates bIDP phase separation by bridging between bIDP chains

To understand the very unusual properties of bIDP-ATP droplets, we carried out all-atom explicit solvent simulations of PM-ATP condensate formation. By following spinodal decomposition ^41^ of a mixture with 16 PM chains and 84 ATP molecules (resulting in charge neutralization), we were able to observe spontaneous formation of a dense slab in a cubic simulation box (Fig. S7 and Supplementary Movie 1), which corresponds to droplet formation in a bulk system. We then extended the simulations in a longer box; the slab is stable in the course of 900 ns of simulations (Fig. 7a, left panel; Fig. S8, first panel). Neither any PM chain nor any ATP molecule dissociates from the slab. These molecules form extensive interaction networks (Fig. 7a, middle panel). ATP molecules bridge between PM chains, with their phosphate groups forming salt bridges with arginine sidechains from multiple chains (Fig. 7a, right panel). Occasionally the riboses form hydrogen bonds with arginines. Interestingly, the adenine bases form both π-π stacking among themselves and cation-π interactions with arginine sidechains (Fig. 7a, right panel and Fig. S9). Frequently these interactions occur together; the resulting multi-layered stacks suggest one mechanism by which PM condensates can accommodate high ATP concentrations.

**Fig. 7.**
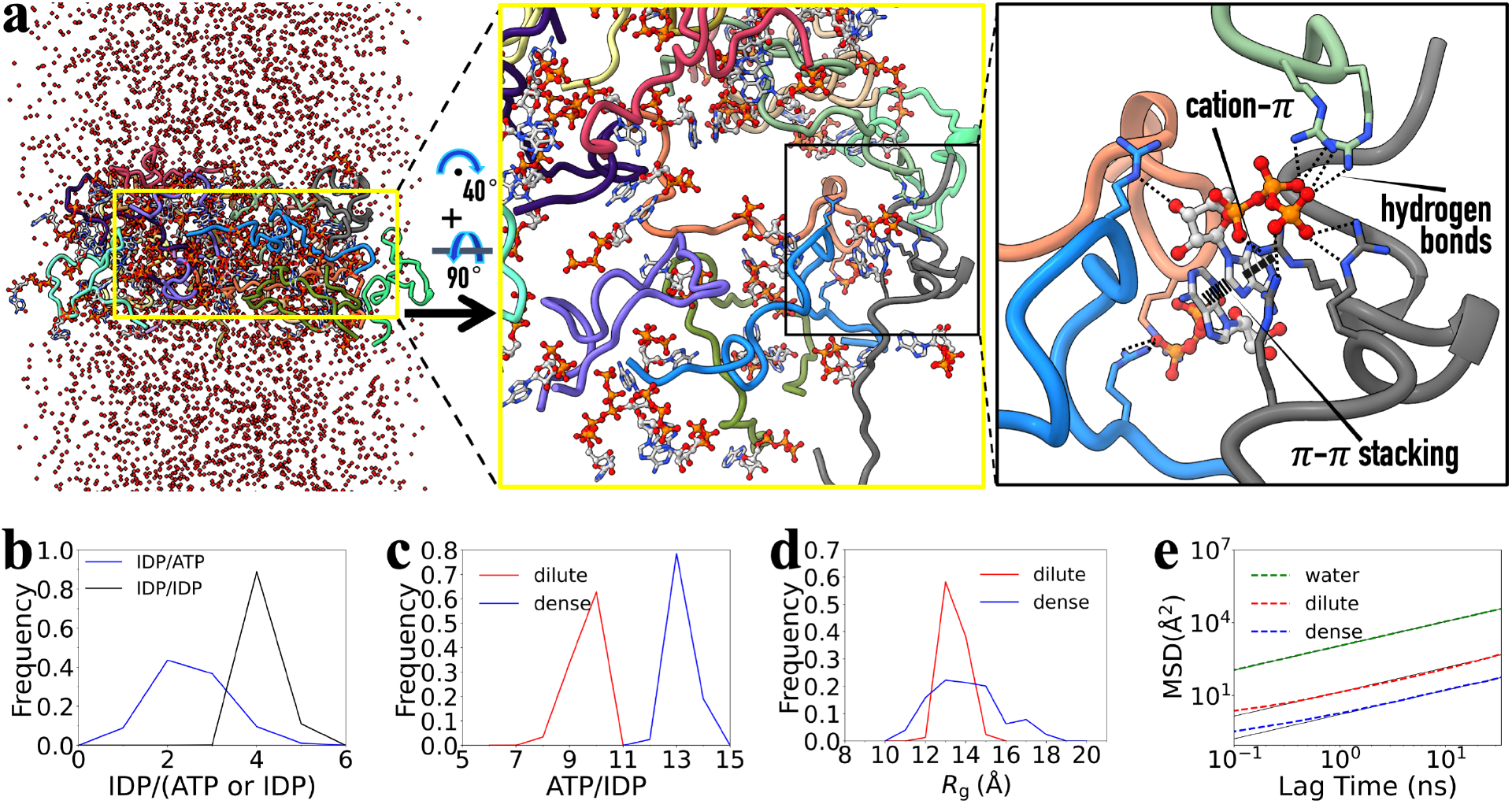
Properties of PM-ATP condensates from MD simulations. (a) Slab formation and intermolecular interactions. In the right panel, arginine sidechains from four PM chains form numerous hydrogen bonds (thin dash) with the phosphates and a ribose on two ATP molecules. One of these arginines also forms a cation-π interaction (thick dash) with an adenine base, which in turn forms π-π stacking (indicated by parallel sticks) with the second adenine, resulting in a three-layer stack. (b) Number of IDP chains in contact with a tagged chain or ATP molecule. (c) Number of ATP molecules in contact with an IDP chain, either in the dense or dilute phase. (d) Distributions of *R*^g^ in the dense or dilute phase. (e) Mean-squared displacements (MSDs) of water and of PM chain in the dilute or dense phase.

In the dense phase represented by the slab, each PM chain on average forms contacts (3.5 Å between heavy atoms) with 4.2 other PM chains, and each ATP molecule on average forms contacts with 2.5 PM chains, highlighting its bridging roles (Fig. 7b). Conversely, each PM chain form contacts with 13.2 ATP chains (Fig. 7c). The crucial role of ATP in stabilizing the dense phase is demonstrated by deformation and dissolution of the slab when ATP molecules are gradually removed (with Cl^−^ ions added for charge neutralization; Fig. S8). To represent the dilute phase, we started simulations of a single PM chain mixed with 10 ATP molecules (with enough Na^+^ ions for neutralization). This single chain on average forms contacts with 9.6 of the 10 ATP molecules (Fig. 7c). We present a snapshot of the single-chain simulations in Fig. S10 to illustrate how these many ATP molecules are accommodated by a single PM chain. In addition to the aforementioned multi-layered stacking, we also observe clustering of phosphates, which are stabilized by arginine sidechains but also mediated by Na^+^ ions. The latter represents another potential mechanism by which PM condensates can accommodate a high concentration of ATP.

The multi-layered stacking and phosphate clustering of the ATP molecules keep the single ATP chain compact, with a radius of gyration (*R*_g_) highly peaked around 13 Å (Fig. 7d). In comparison, the PM chains in the dense phase have a somewhat broader distribution in *R*_g_ with mean ± SD = 14.0 ± 1.6 Å (Fig. 7d). By calculating the mean-squared displacement (MSD) of the chain center of geometry (Fig. 7e), we obtained the translational diffusion constant. In our simulations, the diffusion constant of water molecules is 187 Å^2^/ns. In comparison, the diffusion constants of PM chains are 2.1 Å^2^/ns in the dilute phase and only 0.28 Å^2^/ns in the dense phase. For a tracer molecule with an *R*_g_ about 10 times larger than the size of a water molecule, we expect the diffusion constant to be lower by 10-fold, or 18.7 Å^2^/ns. The 2.1 Å^2^/ns diffusion constant of the single chain must be in large part due to the extensive decoration by the ATP molecules and associated Na^+^ ions. The further slowdown, to 0.28 Å^2^/ns in the dense phase, reflects extensive interaction networks between PM and ATP molecules.

## Discussion

We have shown that ATP mediates the phase separation of two bIDPs. The resulting condensates have very unusual properties, including extraordinarily high ATP concentrations, exceptionally fast fusion speeds, and very low interfacial tension but very high viscosities, thereby manifesting extreme shear thinning. The essential reason for these departures from condensates formed by IDPs alone is that, in those condensates, IDPs chains crosslink to form mesh-like networks (Fig. 8a), whereas in bIDP-ATP condensates, ATP molecules act as bridges between IDPs chains (Figs. 7a and 8b). As a small molecule, ATP is exceptionally facile in forming extensive salt bridges and cation-π interactions with arginine and lysine sidechains of IDPs. Our data show that bIDPs at 1 μM or even lower concentrations can phase separate when mixed with physiological levels (2 -12 mM) of ATP, raising the potential that such phase separation occurs in cells.

**Fig. 8.**
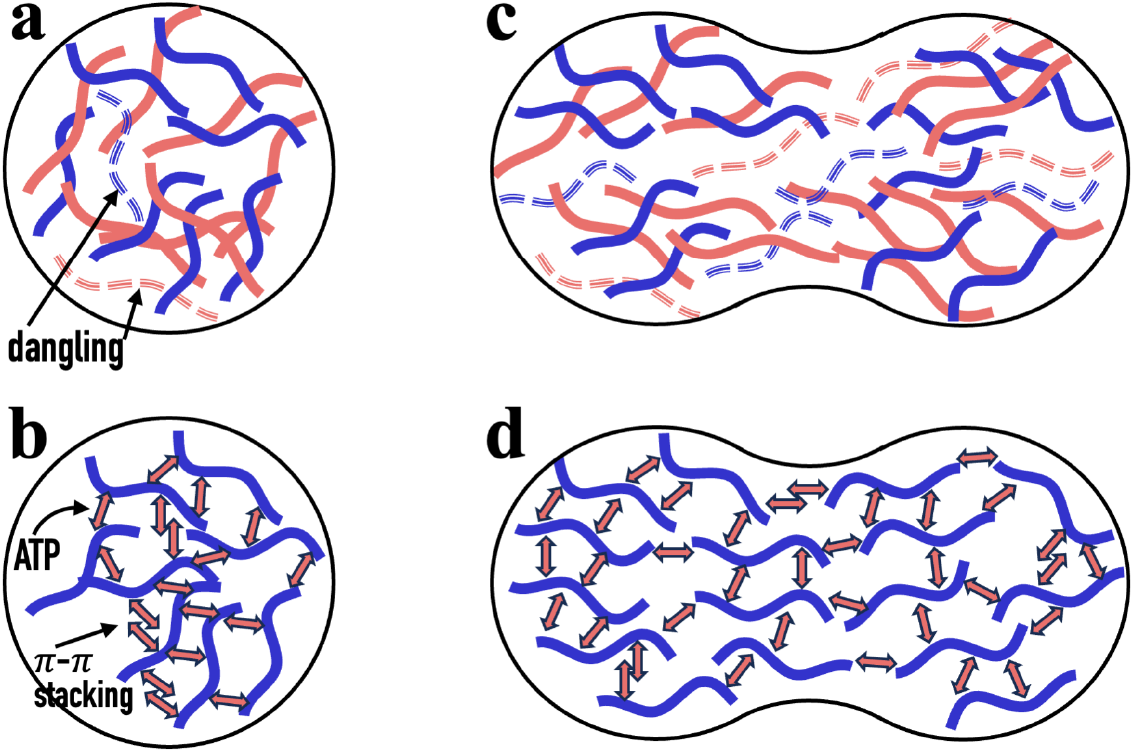
Intermolecular interaction networks in droplets. (a) Droplets formed by oppositely charged IDPs (red and blue). Dangling chains have only a single crosslink with the interaction networks. (b) Droplets formed by a bIDP and ATP. π-π stacking and phosphate clustering allow ATP molecules to bridge between distant chains. (c) Shear thinning in two fusing droplets formed by oppositely charged IDPs, due to chain alignment and chain dissociation. The latter results in more dangling chains. (d) Shear thinning in two fusing bIDP-ATP droplets. Most chains are aligned, allowing them to readily pass each other. ATP bridges quickly reform along the aligned chains.

In previous studies of droplets formed by oppositely charged IDPs, we have seen shear thinning, as measured by the ratio 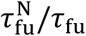, of about 1.2-fold ^40, 42^. For the present pK-ATP droplets, we obtain shear thinning of 25-fold at 5 mM ATP and 67-fold at 10.2 mM ATP. For PM-ATP droplets, the shear thinning is a staggering 108-116 fold. It was suggested previously that shear thinning in IDP droplets occurs due to the alignment of IDP chains along the direction of fusion and dissociation of chains from intermolecular networks (resulting in dangling chains; Fig. 8c) ^40^. For bIDP-ATP droplets, we expect that the IDP chains are even better aligned during fusion, with ATP molecules forming bridges in the orthogonal direction (Fig. 8d). This arrangement allows IDP chains to readily slide past each other, with ATP molecules quickly reforming bridges between them.

It is remarkable that bIDP condensates can soak up so much ATP. Our MD simulations have provided clues for the underlying mechanisms. Salt bridges and cation-π interactions with arginine and lysine sidechains are the primary reason. However, we have also observed that ATP molecules can concentrate around arginine sidechains, where the adenine bases participate in multi-layered stacks and the phosphate groups form cation-mediated clusters. The localized concentration directly strengthens intermolecular interaction networks but also enables bridging between distant chains. These effects explain both the very high zero-shear viscosity (∼15 Pa s) and the long shear relaxation time (∼1000 ms) of PM-ATP condensates. The latter timescale is dictated by the reconfiguration of intermolecular interaction networks. The extensive interactions of a PM chain slow down its dissociation from one subset of ATP molecules prior to reassociation with another subset of ATP molecules. Curiously, at even higher ATP concentrations, the bIDPs form aggregates and fibrils as soon as the samples are mixed, in contrast to the dissolution of condensates reported in many studies. ADP higher than 15 mM does dissolve our bIDP condensates, highlighting the role of the extra phosphate group on ATP.

What may explain the excessively low interface tension of the bIDP-ATP droplets? One possibility is that ATP acts as a “surfactant”, enriched at the interface of the dense and dilute phases. The exposed phosphate groups of these interfacial ATP molecules would have less chances of being stabilized by salt bridges with arginines or lysines than ATP molecules inside a droplet, potentially reducing the interfacial tension. However, the level of interfacial enrichment of ATP cannot be very high; otherwise the resulting repulsion between interfacial ATP molecules would resist droplet fusion, contradicting the rapid fusion that we observed. Our confocal images do not show detectable enrichment of ATP^*^ at droplet surfaces (Fig. 3a, b).

Although PM-ATP and pK-ATP condensates are similar qualitatively, they also exhibit interesting differences. In particular, PM-ATP condensates harbor a much higher level of ATP, and have much higher viscosity and much longer shear relaxation time. These differences can be attributed to much stronger interactions of ATP with arginine than with lysine. PM consists of 64% arginine and no lysine, whereas pK is 100% lysine. The three nitrogen groups of arginine have more ways than the amine of lysine to form salt bridges with the phosphate groups of ATP. The cation-π interaction of the adenine base with arginine is also much stronger than with lysine ^43^. In line with our results, pR-UTP condensates were reported to have substantially higher viscosities than pK-UTP condensates ^16^ (see Supplementary Text 1 for discussion on other related studies).

We observed that ATP and its hydrolysis product ADP produce very different phase diagrams for both PM and pK, recapitulating previous observations on pK-nucleotide mixtures ^25^. As demonstrated in the latter study, enzymatic conversion between ATP and ADP can then be exploited to induce condensation or dissolve condensates. The unusual material properties along with enzymatic control augur well for unique biological functions and potential applications.

## Acknowledgments

We thank Yi Zhang for acquiring TEM images and Fidha Muhammedkutty for contributing scripts.

## Author contributions

Conceptualization: HXZ, DK, RP

Methodology: DK, RP

Investigation: DK, RP, HXZ

Funding acquisition: HXZ

Project administration: HXZ

Supervision: HXZ

Writing: HXZ, DK, RP

## Supplementary information

Materials and methods

Supplementary Figures S1 to S10

Supplementary References

Supplementary Movie 1

