## Supplementary text and figures for "ATP Mediates Phase Separation of Disordered Basic Proteins by Bridging Intermolecular Interaction Networks"

**This PDF file includes:**

Materials and methods
Supplementary Figures S1 to S10
Supplementary references
Captions for Supplementary Movie 1

**Other Supplementary Materials for this manuscript include the following:**

Supplementary Movies 1

### Materials and Methods

#### *Materials*

Protamine (PM) sulfate (catalog # P4020), poly-L-lysine (pK) hydrochloride (15-30 kDa; catalog # P2658), fluorescein sodium salt (catalog # 46960-25G-F), and adenosine 5-diphosphate (catalog # 01905) were from Sigma Aldrich. Adenosine 5-triphosphate disodium salt (catalog # ALX-480-021-G005), Alexa Fluor 647 ATP (ATP*; catalog # A22362) were from Enzo and Thermo Fisher Scientific, respectively. Carboxylate-coated polystyrene beads (2 μm diameter) were from Polysciences Inc. (catalog# 18327-10). For transmission electron microscopy (TEM) imaging, copper grids supported by Formvar/Carbon 300 mesh (product # 01753-F) were from TED PELLA, and phosphotungstic acid (1% aqueous solution; catalog # 19502-1) was from Electron Microscopy Sciences.

#### *Sample preparation*

Imidazole buffer (10 mM, pH 7) was prepared by dissolving 68.08 mg of imidazole (catalog # 396745000, Thermo Scientific) in 100 mL milli Q water and using a magnetic stirrer. pH was adjusted to 7 by adding 3 drops (approximately 70 μL) of 37% (w/w) HCl (catalog # 258148, Sigma Aldrich); the final $Cl^-$ concentration was ~ 8 mM. This "working" buffer was used in all experiments. In some experiments, 150 mM KCl was present. A stock ATP solution at 1 M was prepared in the working buffer, with pH brought to 7 by adding NaOH. ATP dilution was done using the working buffer (estimated $Na^+$ concentration of 24 mM in samples containing 5 mM ATP). ADP solutions were prepared in a similar manner. PM and pK were dialyzed to remove salt before use. In brief, 500 μL of PM (5, 10, or 15 mM) or pK (1.25 mM) stock solution was injected into a dialysis bag (pre-wetted in working buffer) using syringe and placed in 4 L buffer. After overnight on a stirrer, the dialysis bag was taken out and the sample was withdrawn using a syringe. Utmost care was taken to prevent membrane puncture while injecting the sample into or retrieving it from the dialysis bag. All solutions were filtered using 0.22-μm filters (catalog # SLGPR33RS, Millipore Sigma).

Condensates were prepared by sequentially adding the working buffer, KCl (if needed), beads (for interfacial tension and viscosity experiments), PM or pK, and lastly ATP or ADP of desired concentrations in a microcentrifuge tube. All experiments were done at room temperature.

#### *Determination of phase diagrams*

Mixtures of PM or pK with ATP or ADP were observed under an Olympus BX53 brightfield microscope using a 40× objective. Collected images were processed using ImageJ. Phase diagrams were plotted using GraphPad Prism to indicate the observations (homogenous solution, droplets, or aggregates) under the microscope.

#### *Measurement of partition coefficients*

Partition coefficients of ATP* in bIDP-ATP droplets were measured on the confocal module of a LUMICKS dual-trap optical tweezers (OT) instrument. A standard curve was determined on ATP* solutions at concentrations up to 2 mM using confocal scanning with excitation wavelength at 638 nm (0.5% laser power and 0.0128 ms pixel dwell time). Partition coefficients of ATP* were obtained on bIDP-ATP samples prepared with 5 or 10.2 mM ATP plus 1 μM ATP, using the same instrument setting as for the standard curve. PEGylated coverslips [1] were used to produce tall settled droplets for confocal scanning. Confocal images were exported in tiff and processed using ImageJ. The fluorescence intensities in circular regions of interest with a fixed diameter were measured.

To estimate partition coefficients of PM and pK, these bIDPs were first labeled with FITC. In short, fluorescein was dissolved in DMSO to make 10 mM stock. A 500 μL mixture of 25μM pK or 50 μM PM and 400 μM fluorescein was prepared in the working buffer and incubated overnight at room temperature. Excess fluorescein was removed in a PD miniTrap G-25 desalting column (catalog # 28918007, GE Healthcare). A 0.5 μL aliquot of the bIDP-FITC was added to make up a 5 μL bIDP-ATP sample. Confocal scanning was performed with 488 nm excitation (2% laser power and 0.0128 ms pixel dwell time).

***OT-directed droplet fusion***

Two droplets were trapped by the dual-trap OT and grown to equal sizes. The overall laser power was then set to 3% and the split between the traps set to 50%. The droplet trapped by trap 1 was moved toward the droplet trapped by trap 2, and brightfield video recording turned on. Once the droplets were close, the trap-1 droplet was moved in 10-20 nm steps until it came into contact with the trap-2 droplet. Fusion then proceeded spontaneously. The trapping forces were recorded at a sampling rate of 78125 Hz. The time course of the normalized trap-2 force was fit to Eq [1] to obtain the fusion time ($\tau_{\text{fu}}$). The video was analyzed using ImageJ to determine the pre-fusion and post-fusion radii.

***Measurement of interfacial tension by stretching droplets***

Droplet samples were prepared with beads mixed in. A droplet was trapped by both traps (10% overall laser power and 50% split) and grown to a desired size; the traps along with the droplet were then moved to trap two beads one after the other. With trap 2 fixed, trap 1 was pulled away at a slow speed of 0.05 μm/s for 10 s. The sum of the two trapping forces over the extension gave the spring constant $\chi_{\text{sys}}$ of the system comprising trap 1, the droplet, and trap 2 in series [2]. From $\chi_{\text{sys}}$ and the stiffnesses of the two traps, the spring constant $\chi_0$ of the droplet was obtained. Finally the interfacial tension was found as [3]

$$\gamma = \frac{1}{\pi}[\ln(R/a - 1) + 0.68]\chi_0 \qquad \text{[S1]}$$

where $R$ is the radius of the upstretched droplet and $a$ is the radius of the two beads. The droplet radius was obtained from a video recording of the stretching process.

***Measurement of condensate viscoelasticity***

For viscoelasticity measurements, large droplets that were settled on a PEGylated coverslip and had a single bead inside were selected. Trap 1, with 100% of the 10% overall laser power, was used to trap the bead and position it at the center of the droplet. The bead was oscillated over a range of frequencies (0.05 to 40 Hz), with a displacement amplitude of 1 μm (at 0.05 Hz to 2.5 Hz) or 0.5 μm (at higher frequencies). Central positioning of the trapped bead along the $z$ direction was indicated by a phase shift close to 45° between the displacement and trapping force at the lowest frequencies. Displacement and force traces over five oscillation periods were exported as a hdf file and analyzed using a MATLAB program. Briefly, these traces were fit to sinusoidal functions. The resulting ratio between the force and displacement amplitudes and phase shift gave the elastic and viscous moduli $G'(\omega)$ and $G"(\omega)$ [4]. The latter were finally fit to the Jeffreys model of linear viscoelasticity (Eq [3]).

***Imaging aggregates by transmission electron microscope***

On a piece of parafilm, a 20 μL drop of the bIDP-ATP sample, a 20 μL drop of 1% phosphotungstic acid, and three 20 μL drops of milli Q water were placed side by side. A Formvar-coated grid, held by forceps, was placed on top of the sample drop for 2 minutes. The grid was lifted and excess sample was removed by using the ragged torn edge of filter paper to wipe the rim of the grid. The grid was placed on top of the phosphotungstic acid drop for 2 minutes, and then dipped into the water drops one by one to rinse for ~10s each. Extra liquid was again removed by filter paper. Finally the grid was air-dried for 10 minutes in a covered petri dish. TEM images were acquired on a JEM-1400Flash transmission electron microscope equipped with AMT-BIOSPRINT12M-B operating at 80 kV.

***Molecular dynamics simulations***

Using the PM sequence (Uniprot P69014; Fig. 1a), four disordered conformations with radii of gyration close to that expected [5] for an IDP with 33 residues ($R_g = 2.54N^{0.522}$ = 15.8 Å for $N$ = 33) were generated using the TraDES web server [6]. The four PM chains were duplicated into eight copies, modeled by the Amber ff14SB force field [7], and placed randomly in a cubic box with side length of 120 Å and solvated with 52,464 TIP4PD [8] water molecules (Fig. S7a). To neutralize the system, 42 ATP molecules were added in the same water box. Force field parameters for ATP were from Meagher et al. [9].

The system was energy minimized for 10000 steps (2000 steps of steepest descent and 8000 steps of conjugate gradient) using *sander*, followed by a 1 ns equilibration at constant NVT with 1 fs timestep, where temperature was ramped from 0 to 310 K for the initial 100 ps and maintained to 310 K for the remaining 900 ps. Positional restrains were imposed on peptide backbone on each chain with a force constant 5 kcal $mol^{-1}Å^2$. Hereafter the simulation continued with the restraints removed and the timestep increased to 2 fs, first at constant NPT (310 K and 1 bar) for 60 ns and then at constant NVT for 100 ns. At this time the eight PM chains condensed into a glob (Fig. S7b).

We duplicated this glob to build a system with 16 PM chains neutralized by 84 ATP molecules and solvated with 82,412 TIP4PD water molecules, in a cubic box with a side length of 140 Å. To induce the condensation of the 16 PM chains into a slab [10], 70% of the water molecules were removed. The resulting system was energy minimized and equilibrated as described above, except that the last stage was at constant NPT for 630 ns. At this point, a dense slab was well formed (Fig. S7c).

To produce a thicker slab, we removed an additional 10% of the original water molecules. The system was again minimized and equilibrated as above, ending with simulation at constant NPT for 650 ns. The cubic box now had side length of 84.8 Å. The 16 PM chains and 84 ATP molecules from the last frame were solvated in rectangular box, with the length in the $z$ direction elongated 5 times. The number of TIP4PD water molecules was ~94,000, and the total number of atoms was ~387,000. Following energy minimization and equilibration (at constant NPT for 10 ns), two replicate simulations were run at constant NVT for 900 ns. Frames were saved at 100 ps intervals and analysis was done on the last 800 ns.

To ascertain the essential roles of ATP in stabilizing the dense slab, we repeated the simulations in the rectangular box but with portions of the ATP molecules randomly removed. The total number of ATP molecules was reduced to 42, 20, and 15, with $Cl^-$ ions added to provide charge neutralization.

For simulations of a single PM chain, an isolated chain was taken from the simulations with 20 ATP molecules and placed in a cubic box side length of 120 Å, along with 10 randomly positioned ATP molecules. The system was neutralized with 19 $Na^+$ ions and solvated with 54,876 TIP4PD water molecules. The total number of atoms was 220,609. Subsequent simulation steps largely followed those for the rectangular systems, including two replicate simulations at constant NVT for 900 ns. Analysis was done on the last 400 ns.

All the molecular dynamics simulations were run using *pmemd.cuda* [11] in AMBER18 [12]. The cutoff for nonbonded interactions was 10 Å. Long-range electrostatic interactions were treated by the particle mesh Ewald method [13]. Temperature was maintained at 310 K by the Langevin thermostat with a damping constant of 3 $ps^{-1}$. Pressure was maintained at 1 bar by the Berendsen barostat [14]. All bonds linked to hydrogens were constrained using the SHAKE algorithm [15].

CPPTRAJ [16] and in-house python and tcl scripts were used to calculate contacts within 3.5 Å between heavy atoms, radii of gyration, and mean square displacements. For contact calculations, the periodic box was centered at the molecule of interest, to ensure that contacts with other molecules in an image box were all included. ChimeraX [17], Pymol (https://pymol.org), and VMD [18] were used to render images and to make a movie.

**Supplementary Text 1**

Here we comment on a number of previous studies in relation to our findings. Nobeyama et al. [19] determined the phase diagram of pK-ATP mixtures by imaging like us. They focused on very high concentrations of the constituents: pK up to 100 mg/mL (compared to 2.25 mg/mL in our study) and ATP from 50 to 500 mM (compared to our minimum ATP concentration of 0.1 mM). They did not report aggregate formation. One possible reason is that they focused on much higher pK concentrations. Another difference is that their pK chain lengths are ~5 times longer than ours. In addition, we ran our pK through a desalting column to remove salt. Nakashima et al. [20] studied the phase separation of pK (same chain length as ours) mixed with ATP or ADP. At ~40 μM pK, the threshold ATP concentration was ~ 1 mM, the same as ours (Fig. 2b), though their mixture contained 5 mM $MgCl_2$. Their maximum ATP was 15 mM, and no aggregate formation was reported. For ADP, the threshold concentration was ~2 mM, compared with ours at 1 mM (Fig. 2d). At 15 mM ADT, their turbidity data did not show a decrease to 0, as we would predict from our phase diagram (Fig. 2d). However, this point is near the phase boundary, which may be subject to effects such as the 5 mM $MgCl_2$ in their samples.

There were two reports of phase separation for PM mixed with polyvalent anions [21, 22]. In one, the anion was sulfate; in the other, the anions were triphosphate and citrate. In the latter study, Kim et al. also reported material properties of PM-citrate condensates. Using confocal imaging, they obtained an inverse fusion speed of ~10 ms/μm, comparable to our value for PM-ATP condensates (Fig. 4f). Using particle tracking, they obtained a viscosity of 0.07 Pa s, which is ~200-fold lower than our value. Our viscosities for pK-ATP and PM-ATP condensates are comparable to those reported for pK-UTP and pR-UTP condensates by Fisher et al. [23], also using particle tracking. Kim et al. went on to deduce the interfacial tension using Eq [2], which is only valid for Newtonian fluids. Our data show that PM-ATP condensates deviate enormously from Newtonian fluids and exhibit extreme shear thinning, and hence Eq [2] is totally unusable.

Williams et al. [24] studied the coacervation between a basic polymer, poly(diallyldimethylammonium) chloride, and ATP. In these coacervates, the partition coefficient of ATP was of the order of 15, much lower than what we measured for PM-ATP and pK-ATP droplets. The viscoelastic properties were measured by a viscometer. The zero-shear viscosity was 0.9 Pa s, similar to our value for pK-ATP condensates; the viscous and elastic moduli also look similar to ours. Interestingly, their condensates were probed by wide angle X-ray scattering, showing the absence of a band corresponding to ATP π-π stacking, in contrast to what we observed in MD simulations of PM-ATP condensates.

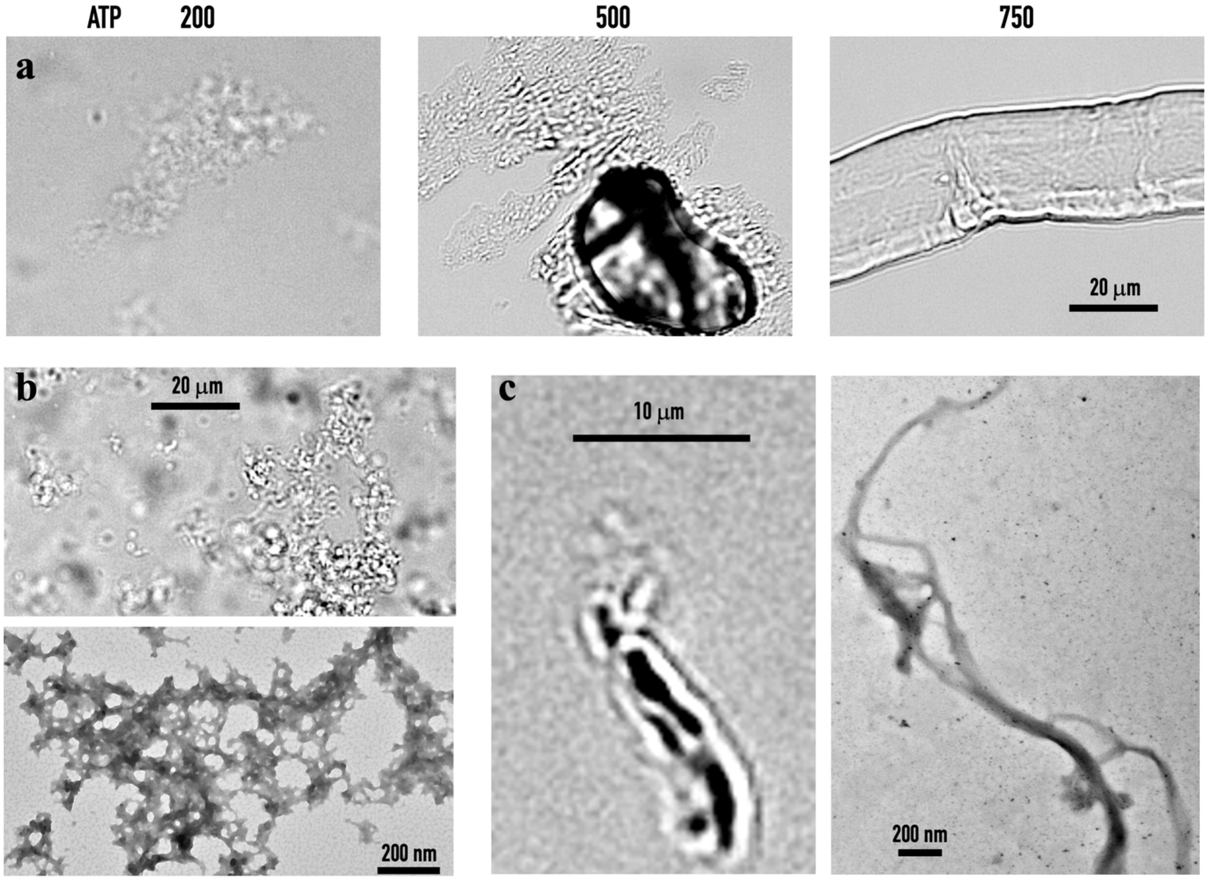


**Supplementary Fig. S1.** Aggregate formation when bIDPs are mixed with high concentrations of ATP. (a) Brightfield images showing that 10 μM PM forms amorphous aggregates when mixed with 200 mM ATP, fibril-like aggregates with 500 mM ATP, and fibrils with 750 mM ATP. (b) A mixture of 50 μM pK and 100 mM ATP in the presence of 150 mM KCl forms amorphous aggregates, as shown by both brightfield (top panel) and TEM (bottom panel) images. (c) A mixture of 1 μM pK and 100 mM ATP without KCl forms fibril-like aggregates, as shown by both brightfield (left panel) and TEM (right panel) images.

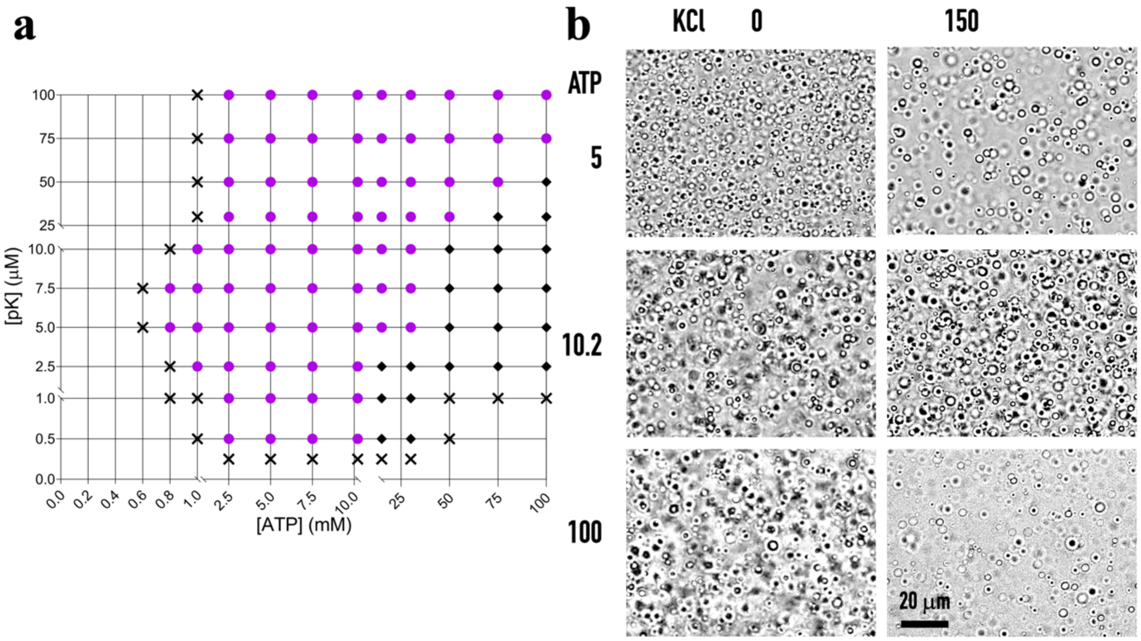


**Supplementary Fig. 2.** Effects of 150 mM KCl on the phase diagram of pK-ATP mixtures. (a) Phase diagram in the presence of 150 mM KCl. Magenta circles indicate droplet formation; black diamonds indicate aggregate formation; crosses indicate homogeneous solution. (b) Brightfield images of droplets formed by mixing 100 μM pK with ATP at the indicated concentration (in mM), without or with 150 mM KCl.

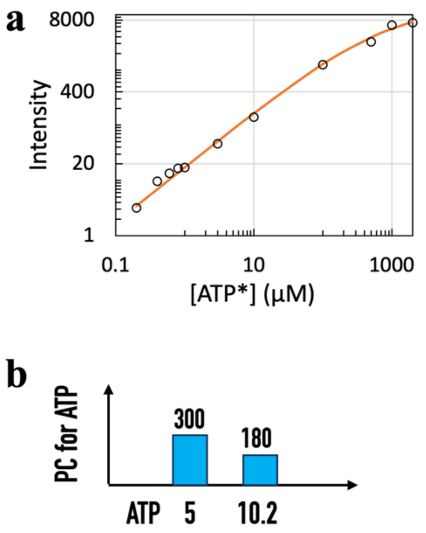


**Supplementary Fig. 3.** Partition of ATP inside droplets by confocal microscopy. (a) Standard curve for ATP*. (b) Partition coefficients (PCs) of ATP* in droplets formed by mixing 100 µM pK and 5 or 10.2 mM ATP in the presence of 150 mM KCl.

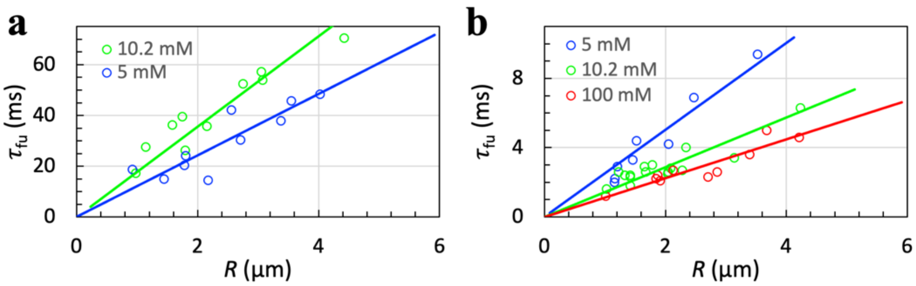


**c**

| | PM | | pK | | |
|---|---|---|---|---|---|
| ATP (mM) | 5 | 10.2 | 5 | 10.2 | 100 |
| $\tau_{fu}/R$ (ms/μm) | 12.1±0.8 | 17.8±0.8 | 2.51±0.13 | 1.44±0.06 | 1.12±0.05 |

**Supplementary Fig. 4.** Fusion speeds from directed droplet fusion by optical tweezers. (a) Fusion time as a function of initial droplet radius, for droplets formed by mixing 100 μM PM with 5 or 10.2 mM ATP in the presence of 150 mM KCl. (b) Corresponding results for pK-ATP droplets. (c) Inverse fusion speeds of bIDP-ATP droplets in the presence of 150 mM KCl.

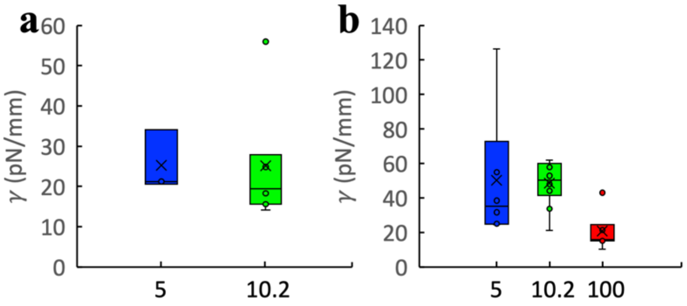


c

| | PM | | pK | | |
|---|---|---|---|---|---|
| ATP (mM) | 5 | 10.2 | 5 | 10.2 | 100 |
| γ (pN/µm) | 25±6 | 26±14 | 50±36 | 49±12 | 21±10 |

**Supplementary Fig. 5.** Interfacial tension from stretching droplets by two optically trapped beads. (a) Interfacial tension of PM-ATP droplets formed by mixing 100 µM PM with ATP at the indicated concentration (in mM) in the presence of 150 mM KCl. (b) Corresponding results for pK-ATP droplets. (c) Mean values of interfacial tension in the presence of 150 mM KCl.

**a**

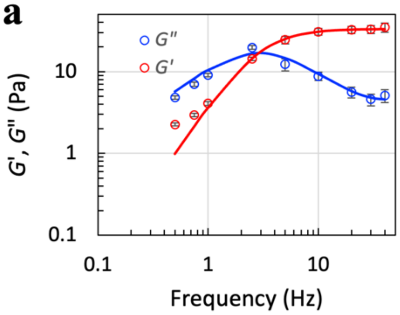


**b**

| | pK | |
|---|---|---|
| ATP (mM) | 5 | 10.2 |
| $\eta_0$ (Pa s) | 0.029 | 0.014 |
| $\eta_1$ (Pa s) | 0.62 | 0.88 |
| $\tau_1$ (ms) | 27 | 34 |

**Supplementary Fig. 6.** Viscoelasticity measured by oscillating a trapped bead inside droplets. (a) Elastic and viscous moduli of pK-ATP droplets (100 μM pK and 5 mM ATP with 150 mM KCl). (b) Viscosities and shear relaxation times of pK-ATP droplets in the presence of 150 mM KCl.

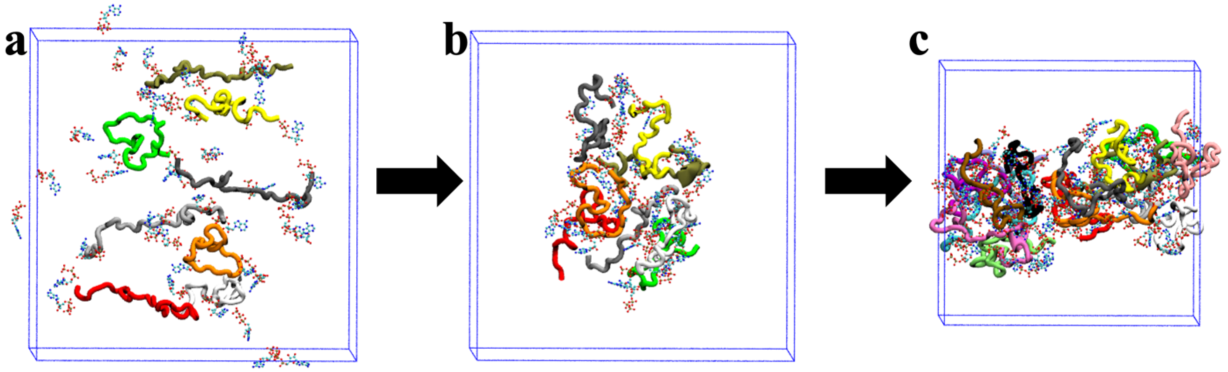


**Supplementary Fig. 7.** A series of simulations leading to the formation of a dense slab. (a) The initial configuration of 8 PM chains and 42 ATP molecules, dispersed in a cubic box with 120 Å side length. (b) A dense glob formed after 160 ns of simulation. (c) That glob was duplicated and solvated in a cubic box with 140 Å side length. After 70% water removal, the box shrunk and the 16 PM chains and 84 ATP molecules condense into a slab after 630 ns of simulation.

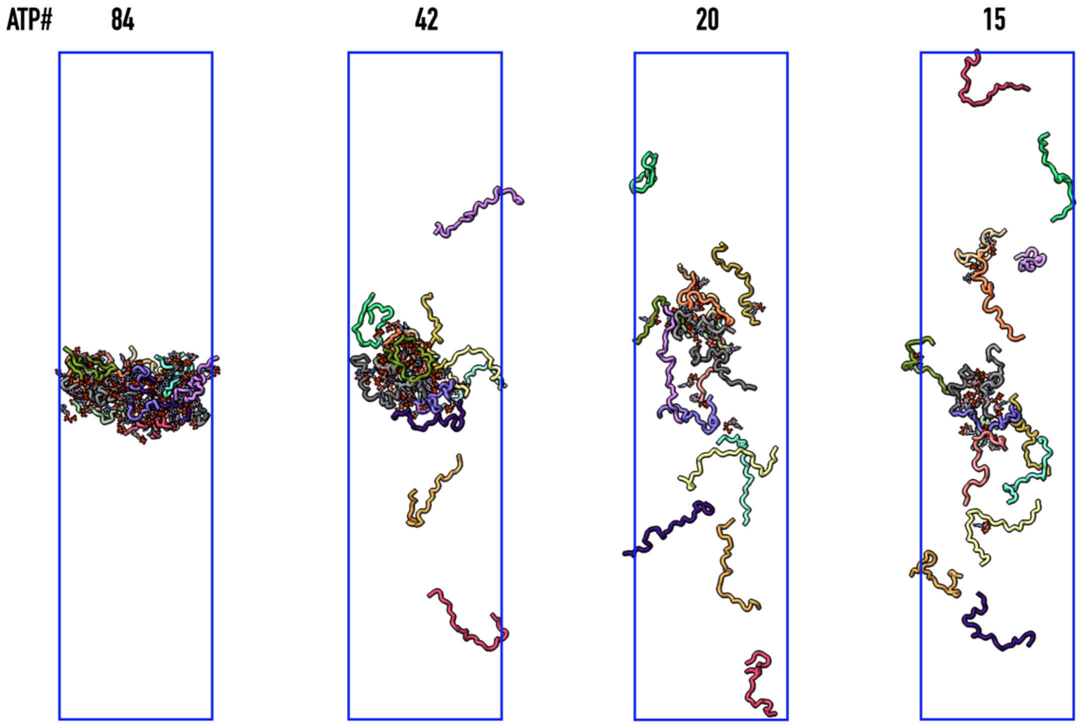


**Supplementary Fig. 8.** ATP maintains the stability of a dense slab. The dense slab is stable when 84 ATP molecules are present to neutralize the charges on the 16 PM chains, but starts to deform and eventually dissolve when ATP is removed. The frame of the 84-ATP system is at 400 ns of a 900-ns simulation (same as shown in Fig. 7a); the frames for the systems with reduced ATP numbers are at 201 ns.

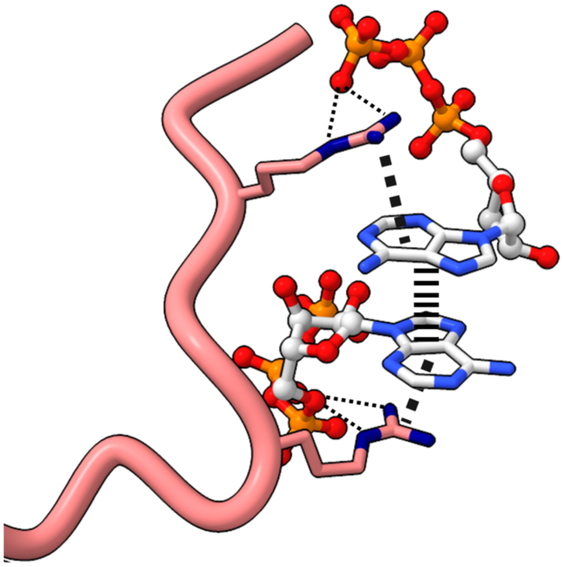

**Supplementary Fig. 9.** Cation-π interactions between arginine sidechains and ATP adenine bases, indicated by thick dash, in the dense slab shown in Fig. 7a. These cation-π interactions are part of a four-layer stack, involving two π-π stacked (indicated by parallel sticks) adenines in the middle and two arginines on the opposite sides. In addition, the arginine sidechains also form salt bridges with ATP phosphate groups (thin dash).

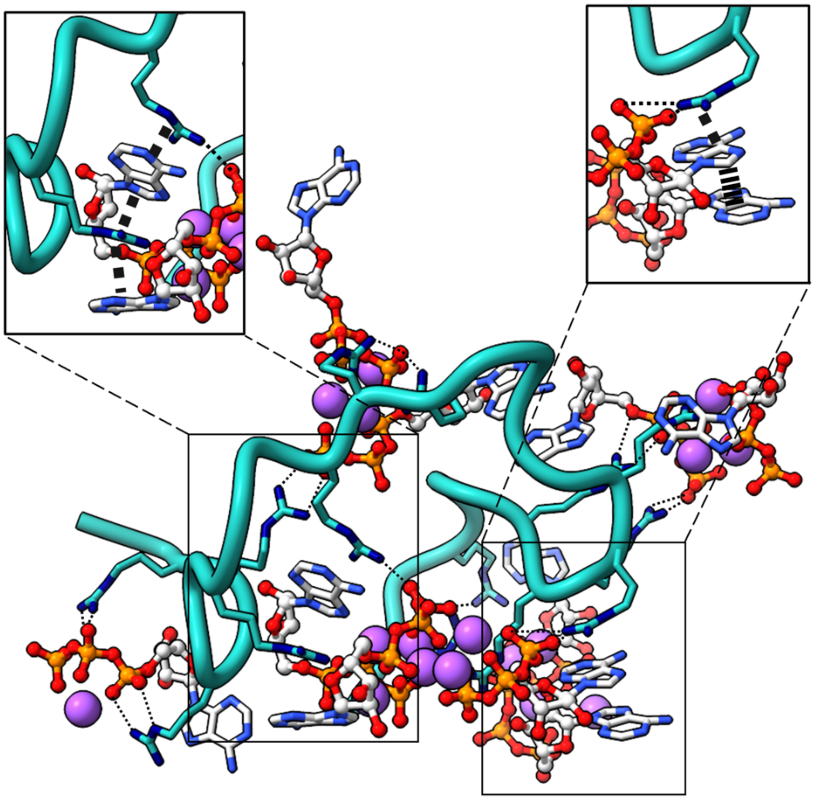

**Supplementary Fig. 10.** A snapshot from MD simulations of a single PM chain in the presence of 10 ATP molecules. The dense population of ATP molecules results from salt bridges (thin dash) and cation-π interactions (thick dash) with arginines, π-π stacking, and clustering of phosphate groups around $Na^+$ ions (magenta spheres). The zoomed view on the top left shows a four-layer stack involving two arginines intercalating two adenines; the zoomed view on the top right shows a three-layer stack involving an arginine and two π-π stacked adenines (indicated by parallel sticks); these arginines also form salt bridges with phosphate groups.

**Supplementary Movie 1.** A series of simulations leading to the formation of a dense slab comprising 16 protamine chains and 84 ATP molecules. Three snapshots are shown in Fig. S7, with further details given in the caption of that figure.
